# Easyreporting simplifies the implementation of Reproducible Research Layers in R software

**DOI:** 10.1101/2020.12.07.414417

**Authors:** Dario Righelli, Claudia Angelini

**Affiliations:** Department of Statistical Sciences, University of Padova, Padua, Italy; Istituto per le Applicazioni del Calcolo “Mauro Picone”, National Research Council, Naples, Italy

## Abstract

During last years “irreproducibility” became a general problem in omics data analysis due to the use of sophisticated and poorly described computational procedures. For avoiding misleading results, it is necessary to inspect and reproduce the entire data analysis as a unified product. Reproducible Research (RR) provides general guidelines for public access to the analytic data and related analysis code combined with natural language documentation, allowing third-parties to reproduce the findings. We developed *easyreporting*, a novel R/Bioconductor package, to facilitate the implementation of an RR layer inside reports/tools without requiring any knowledge of the R Markdown language. We describe the main functionalities and illustrate how to create an analysis report using a typical case study concerning the analysis of RNA-seq data. Then, we also show how to trace R functions automatically. Thanks to this latter feature, *easyreporting* results beneficial for developers to implement procedures that automatically keep track of the analysis steps within Graphical User Interfaces (GUIs). *Easyreporting* can be useful in supporting the reproducibility of any data analysis project and the implementation of GUIs. It turns out to be very helpful in bioinformatics, where the complexity of the analyses makes it extremely difficult to trace all the steps and parameters used in the study.

## Introduction

Due to accidental mistakes or misusage of sophisticated computational methods, many research findings in omics science are considered false (or partially incorrect) [1]. Moreover, in several cases, published results are not entirely reproducible due to the lack of information. For example, the analysis of the massive amount of omics data produced by high-throughput technologies requires combining several different methodologies from the preprocessing, data cleaning, and normalization to the downstream analysis. Therefore, it becomes challenging to trace all the steps and the parameters used within a complete analysis. Consequently, the lack of details, such as user-parameter or subtle data manipulation made with small code lines not reported in the material and methods sections of a manuscript, can lead to findings that are not reproducible. To prevent misleading results, several authors suggested adopting some best practices [2–4] that should help in publishing reproducible results. Nevertheless, the proposed approaches can be time-consuming and require significant effort by researchers. Therefore, to fully exploit the advantages of Reproducible Research (RR), it is still necessary to provide tools that can trace all the details using automatic procedures [5, 6].

Recently, the scientific community proposed several approaches to support RR by developing tools that require a lower cost in terms of time and efforts to be used [5–9]. Among the different approaches, one common idea is to describe the steps with an analysis report built up as a mixture of natural language sentences along with computational language and graphical outputs. This document should include: i) the analyzed data, ii) the Code Chunks (CCs), iii) results and intermediate outputs (as tables and figures), and iv) all information that can enhance the work comprehensibility and reproducibility. Using human-readable reports instead of other procedures, such as dockers, has the advantage that the final document can be easily understood by non-expert users, whereas dockers require computationally experienced users. Moreover, a human-readable report can be enriched with comments and favors knowledge transfer. Nevertheless, the two approaches are complementary and can be combined to achieve full reproducibility in terms of input/output of each algorithm/function and the possibility to re-create a computational environment that does not depend on specific user installations.

The R community proposed several solutions based on the literate statistical programming, like *sweave* [10], *knitr* and *rmarkdown* [11]. Within this framework, the authors can release a data analysis as a human-readable document that incorporates data, computational methods (including the short lines of code that are often omitted in a high-level description of the computational procedure), user-parameters, tables, and figures. Moreover, this report is automatically updated each time the analyst introduces some workflow changes to preserve complete reproducibility. R Studio (https://rstudio.com) already contains several functionalities that can help an analyst compiling detailed reports. Other R packages, such as Drake [12], go through the same directions.

Even though *rmarkdown* is very popular inside the R community, its usability when developing automated tools as Graphical User Interfaces (GUI) or packages is limited. Despite several efforts, incorporating a RR layer in other software and automatically tracing all the steps performed during a point-and-click analysis is still challenging. In the past, we proposed a solution with the RNASeqGUI [13, 14] project. RNASeqGUI is a GUI for analyzing RNA-seq data that automatically traces the analysis steps and reports them in a unique report. Although very useful, this solution did not allow the user to add personal comments, a particularly relevant requisite for knowledge transfer. Moreover, its implementation was time-consuming.

In light of these reasons, we developed *easyreporting*, a R/Bioconductor package allowing us to construct HTML reports that automatically incorporate comments with data, code, plots, and tables. In this work, we describe the *easyreporting* class and its methods. Then, we show i) how *easyreporting* can be used to generate user analysis reports and ii) how *easyreporting* can be used to implement GUIs that automatically trace their functions and produce an analysis report.

## Materials and methods

### Implementation

*Easyreporting* is an open-source R/Bioconductor package aimed to 1) support analysts to speed up the compilation of their analysis reports (without learning the *rmarkdown* language) and 2) help developers to integrate a RR layer inside their software products (such as GUI). While the first aim can also be achieved using other similar tools, the latter constitutes one of the main advantages of our solution. In such a way, thanks to minimal efforts on the developers’ side, the end-user (that can be not necessarily aware of the R or the *rmarkdown* language) can obtain an *rmarkdown* file that incorporates the source code generated during the analysis with the user-friendly tools. Once compiled, this document can then be published as supplementary material of a scientific article, helping the interested community to reproduce the computational part of the work entirely, as suggested in [15]. Moreover, the document can be easily organized into sections, describing different analysis steps, and enriched with natural language comments, making the report more explainable to increase the knowledge transfer.

### General Description and Initialization

*Easyreporting* is structured as an S4 class representing a schematic view of a *rmarkdown* file (see Fig 1). Thanks to *easyreporting*, an analysis report can be seen as a particular instance of the package class, where the attributes represent the report characteristics. Within this class, the available methods are useful for attribute manipulations and for inserting comments and organizing section titles inside the report.

**Fig 1.**
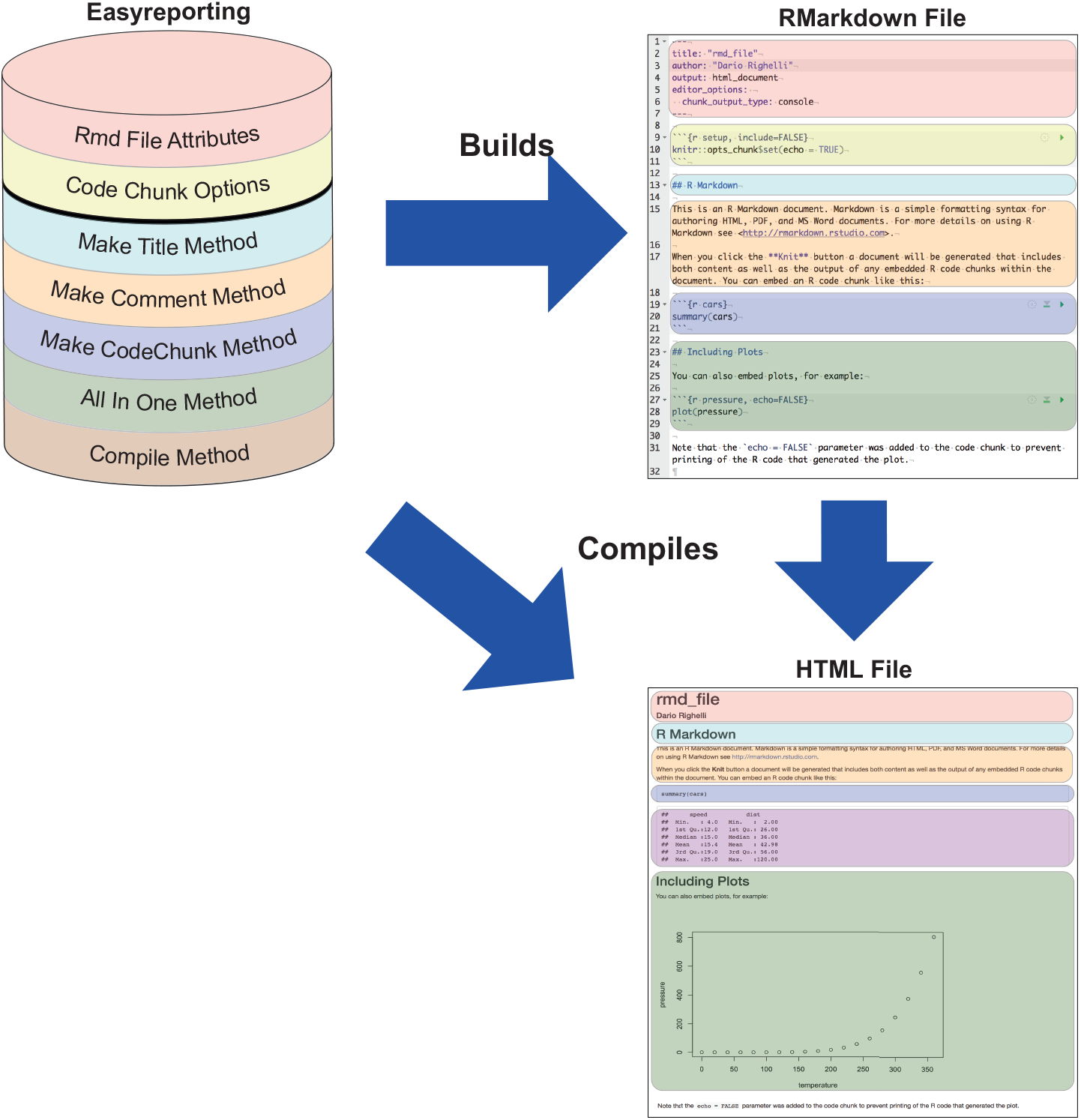
The *easyreporting* class package is a representation of a *rmarkdown* file. The color codes indicate which attribute/method represents the same-color portion of the *rmarkdown* first, and the compiled final HTML report then.

When *easyreporting* is used to create a report for novel analysis, the analyst needs to initialize an instance of the *easyreporting* class with the *easyreporting()* constructor function, passing as mandatory arguments the path and the name of the report file accompanied by its title. Optionally, it is possible to specify an author. In this way, each analysis/project is uniquely associated with a specific *easyreporting* instance, and hence to the corresponding *rmarkdown* file. The initialization step can be transparent to any GUI user since the developers handle the tool’s backend.

During the initialization, the class constructor automatically creates the report file inside the specified folder tree, setting up its header and declaring the general options for the *rmarkdown* file. As soon as the analyst or the user proceeds with his analysis, the *rmarkdown* file is updated with a new CC each time the analysis software performs a new analysis step. When the analysis is complete, it is possible to compile the report using the *compile* method, which produces the final report (in HTML format) and appends a final CC with the *sessionInfo* to trace all the packages versions used for the analysis.

### General Exploitation

The *easyreporting* class is equipped with several methods for *rmarkdown* CC construction (see Table 1 for the full list). Once an *easyreporting* instance is available, it is possible to organize the report by inserting up to six levels of titles by using the *mkdTitle* method. It is also possible to add natural language comments with *mkdGeneralMsg*. The latter feature is particularly relevant to make the analysis more understandable.

**Table 1.** Attributes and Methods of the *EasyReporting* class.

| Attributes | Description |
| --- | --- |
| filenamePath | the report file name with the absolute path |
| title | the title of the report |
| author | the auhor |
| documentType | actually this is set to HTML |
| optionList | a list of R Markdown options |
| Methods | Description |
| mkdTitle | Inserts an R Markdown title inside the report |
| mkdGeneralMsg | appends a general message to the report |
| mkdGeneralTitledMsg | appends a title and a general message to the report |
| mkdVariableAssignment | includes a variable assignment into the report |
| mkdCodeChunkSt | creates a CC start |
| mkdCodeChunkEnd | creates a CC end |
| mkdCodeChunkComplete | creates a complete CC |
| mkdCodeChunkCommented | creates a complete CC with a previous comment |
| mkdCodeChunkTitledCommented | creates a complete CC with a previous comment and a t |
| mkdSourceFiles | includes a list of source files inside the CC |
| compile | prints sessionInfo and compiles the R Markdown file |
| setOptionsList | set an optionList to the class |
| getOptionsList | returns the optionList from the class |
| getReportFilename | returns the report filename with path |

For the implementation of the CCs creation, we suggest two main approaches based on the methods available in the class (see the examples in Listings 2 and 3 shown in the Results Section): i) The first approach builds a CC as a typical step-by-step process. It consists of opening a CC (*mkdCodeChunkSt*), adding variables assignments and/or function callings (*mkdVariableAssignement*), and finally closing the CC (*mkdCodeChunkEnd*). In this approach, it is possible to add comments with *mkdGeneralMsg* before closing the CC. ii) The second approach builds the CC in a single step by using the *mkdCodeChunkComplete* method. The method automatically embraces the tracking code into a new CC, while the user has to take care of the variable assignments and/or the function he/she wants to trace by passing it as an argument. In this approach, the user can also add personal comments passing them as an additional argument. Then, *easyreporting* automatically adds the comment before the new CC.

The first approach appears useful when we need to carry out several R commands in a single CC. It is similar in the spirit to the functionalities offered by R-studio or other development environments, however since it is entirely command-line, it can be easily used on systems with limited development capabilities. By contrast, the second approach is more appropriate for tracing a single function call automatically. In the next section, we will show how this second possibility can be particularly useful to wrap functions performing a specific step and trace their execution within GUIs.

### Implementing automatically tracing functions and their usage within GUIs

The previously described CCs creation approaches can be adapted to trace several steps of an analysis pipeline and end-up with a nicely formatted and detailed analysis report. However, they require the analysts to manually trace each step of the analysis (as he could also do with other available tools). Consequently, the above approaches are useful for generating analysis reports (that was the first aim of the *easyreporting* package). However, they are not suited for the automatic tracing of the steps of an analysis performed using point-and-click approaches thorough GUIs (the second aim of the *easyreporting* package).

In the last decade, GUIs are becoming very popular in bioinformatics because they simplify computational analysis allowing non-expert users to choose among several computational procedures, algorithms, and parameter settings, see for example [13, 14, 16–18]. In particular, the *shiny* (https://shiny.rstudio.com) libraries simplified the development of GUIs that incorporate the power of the statistical R language and the wide-amount of open-source packages available in repositories such as Bioconductor (https://www.bioconductor.org).

Nevertheless, computational studies obtained from GUIs might lack reproducibility since tracking all user choices is still challenging. To face this limit, the developers have to implement a RR layer when designing the GUI’s back-end so that the final users can benefit from a better quality product. Moreover, the RR layer has to be transparent but understandable to not-expert users. Ideally speaking, at the end of the analysis, the user should have a human-readable report analogous to the one obtained using command-line approaches.

*Easyreporting* methods can be easily adapted to support the automatic tracing of any given function by combining a rendering function that performs the required step with a wrapping function that traces its execution. The wrapper function (WF) needs an *easyreporting* instance, and the arguments of the function to be traced (TF). Then, the developer inserts the WF in the back-end of the interface (i.e., the server if the context of a GUI implemented with the shiny library) in the TF place. The front-end of the interface (i.e., the UI with the shiny library) remains unchanged. When the user interacts with the interface to invoke the TF, the back-end will invoke the WF, which will call both the TF function of interest and trace its usage with all parameters. In brief, employing wrapper functions makes it possible to implement a reproducible research layer within the GUI without implementing all the tracing *rmarkdown* code. Listings 4-6 illustrate a specific case with a volcano plot, and Figure 2 schematically represents the entire workflow of information.

**Fig 2.**
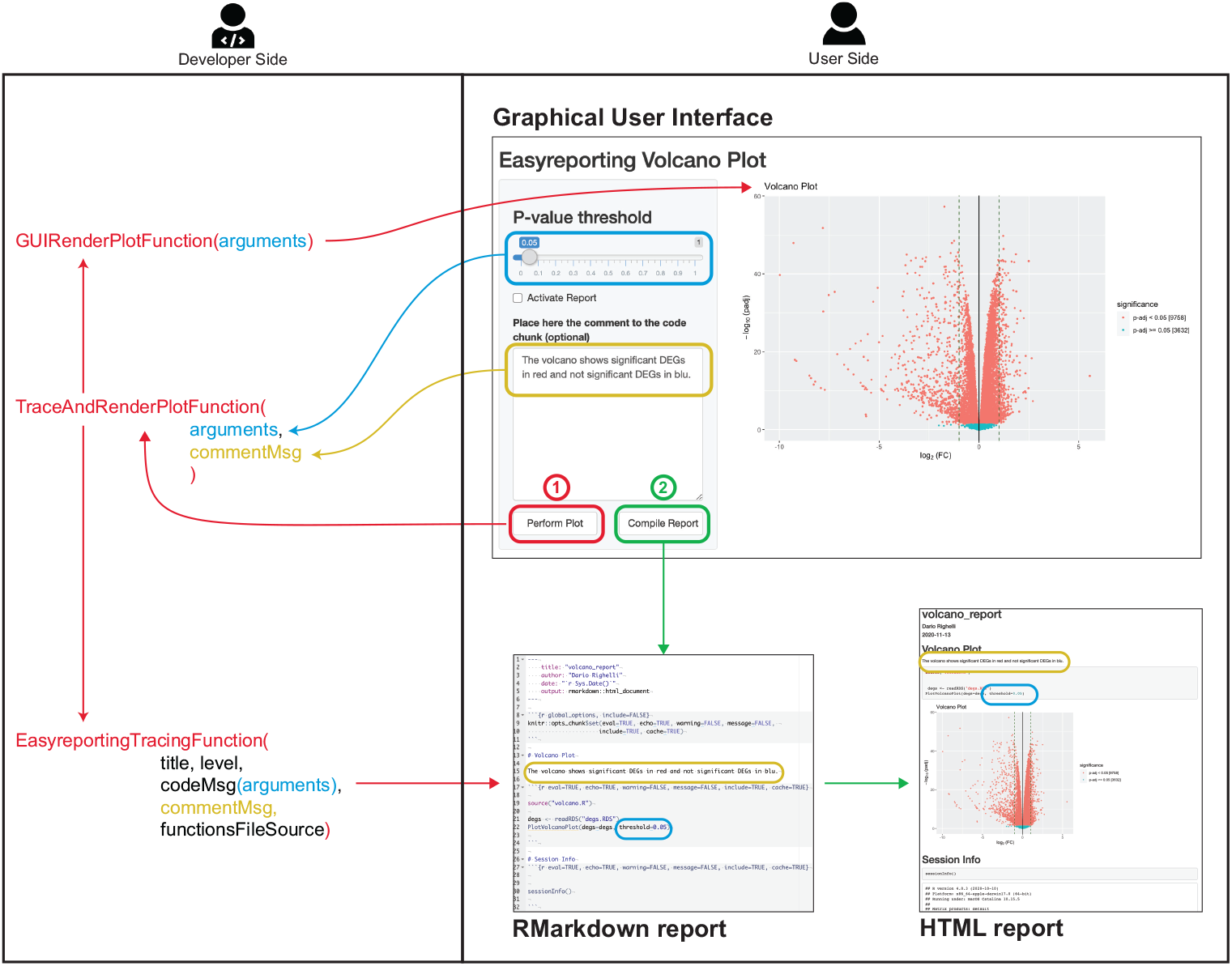
Example of a Graphical User Interface working with *easyreporting* package. The right panel shows the user side of the software with the user interactive GUI. The left panel shows the RR layer transparent to the user, which traces the performed step as organized by the developer. In this example, the GUI allows performing a Volcano Plot starting from a DEGs data frame that includes Log Fold change and p-values as column variables. With the p-value threshold selector (in blue), the user can set the threshold for the significance of the p-values. Moreover, the user can add a personal comment (in yellow) that wants to add alongside the traced code. The user has to press the perform button (in red) to visualize the plot into the GUI. In the meantime, the back end will trace the step into the *rmarkdown* file and will return the plot to the user interface. The user can then compile the report through the compile report button (in green) and visualize the HTML report. The back-end will execute the *compile()* function and return the result to the interface. To make this possible, the developer needs to include a RR layer into the GUI engine (i.e., the shiny server-side). For example, the developer can define a traced function (TF) that renders the Volcano-plot and a wrapper function (WF) that takes as input the input arguments from the graphical GUI (i.e., the shiny UI side). The WF code can be structured to render the plot into the graphical GUI and make a call to an *easyreporting* method, combining the input arguments with the easyreporting class method required ones, to write a new CC into the analysis rmarkdown. Additionally, the developer has to bind the *Compile Report* button with the *compile easyreporting* method to generate the HTML final report.

## Results

To illustrate the capabilities of *easyreporting* we first show its usage for generating an analysis report in a case study concerning the analysis of RNA-seq data (see Supplementary File 1 for details), then we illustrate how to implement a simple GUI that automatically traces its usage and the choice of parameters to produce a report.

### Easyreporting for the creation of analysis report

The RNA-seq data used in the example allows investigating the differences in CD8+ dendritic T-cells of the immune response of two different antibodies compared with control, see [19] for more details. We chose this illustrative example since it is well-known that the analysis of RNA-seq data can lack reproducibility [20]. The dataset contains the raw counts of 37991 genes and is composed of two replicates for each of the three conditions: DEC (fd-scaDEC-205 antibody samples); E2 (E2 antibody samples) and UNTR (control samples). For illustrative purposes, in our Supplementary file 1, we start the analysis from the raw count-matrix. Moreover, we released the raw counts as supplementary data with the *easyreporting* package, allowing the readers to reproduce our example. The naive pipeline will first load the data, perform some diagnostic plots, filter and normalize the raw counts, and visualize the principal component projection. It will then perform differential gene expression analysis and depict the results as a Venn diagram and MA-plots. A specific CC describes each phase.

In the following, we show the main fundamental steps that a user can adapt to any analysis, and we refer to the Supplementary File 1 for the detailed description of the remaining steps and to the Supplementary File 2 for the complete report.

### Report Initialization

After loading the *easyreporting* package in the R environment, the analyst needs to initialize an analysis report by providing the file name (i.e., “rnaseq report”) and the title of the document (i.e., “RNA-seq Analysis Report”). It is also possible to specify an author (i.e., “Dario Righelli”). For simplicity, we set-up a project directory path starting from the working directory for our report, but the user can choose other locations by setting the *filenamepath* parameter. The initialization is carried out by using the *easyreporting()* function. Note that the *filenamepath* and *title* are mandatory parameters, while the author is optional. The following Listing 1 code illustrates the initialization of a report.

**Listing 1.**
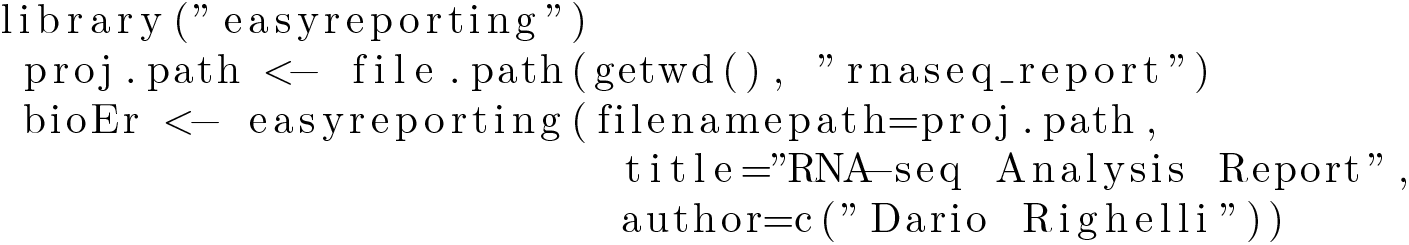
Initialization chunk

### Creation of a chunk of code

Once the analyst has initialized the report, he/she can add a CC for each step of the analysis. As mentioned in the General Exploitation section, EasyReporting provides two main approaches for adding CCs within a report: 1) building up the CC step by step (as shown in Listing 2) and 2) using several kinds of wrapper functions (as shown in Listing 3).

As mentioned above, in the first case, the analyst has to use the *mkdCodeChunkSt* to open a new CC. Then, he/she needs to add the code to markdown, by using the *mkdVariableAssignment* and/or the *mkdGeneralMsg* functions, for tracking variables and functions. Finally, the analyst has to close the CC using the *mkdCodeChunkEnd* function. The following Listing 2 code illustrates a step-by-step CC for loading the counts matrix released with the package.

**Listing 2.**
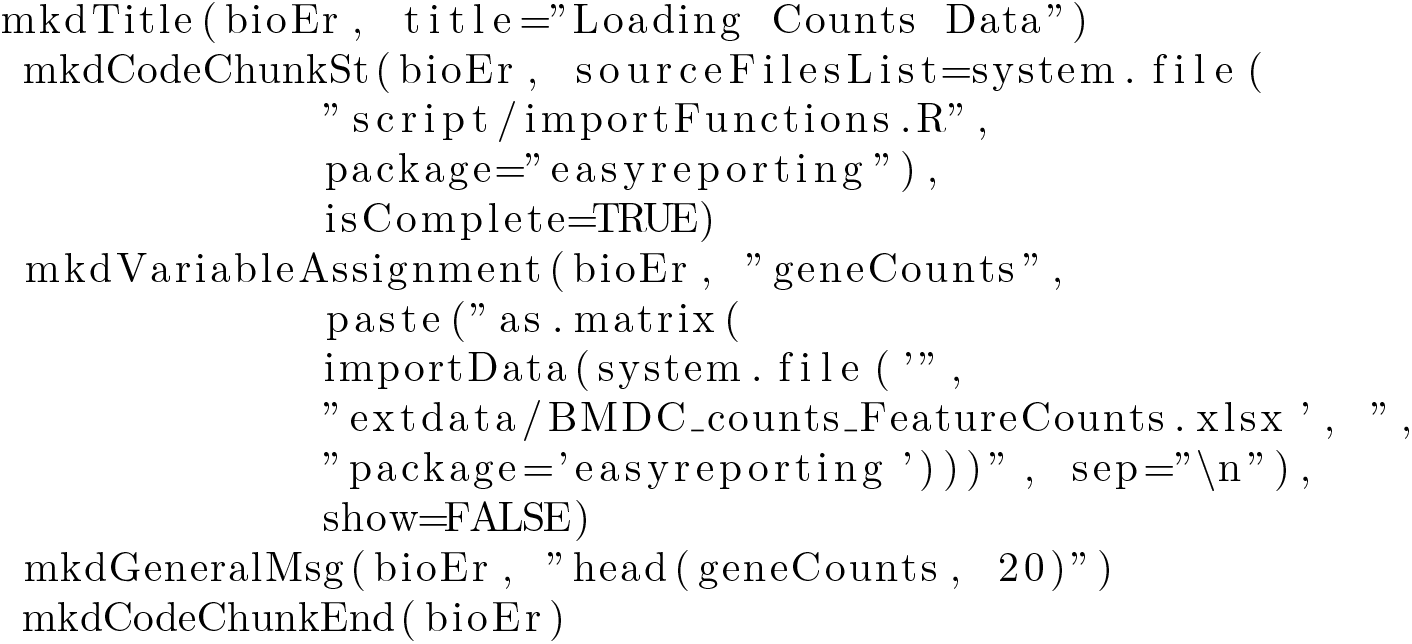
Step-by-step chunk construction

Although the first approach leaves complete freedom to the analyst, it can be tricky for small CCs. The second approach can be more straightforward for small CCs. To this purpose, the *mkdCodeChunkComplete* function allows tracing the steps through the message parameter. The following Listing 3 code illustrates an example of a single step CC. As for the above CC, we assume that the analyst wants to read the raw counts using a user-defined function, here named “importData.R”, that we stored into the importFunctions.R file available in the package “script” folder.

**Listing 3.**
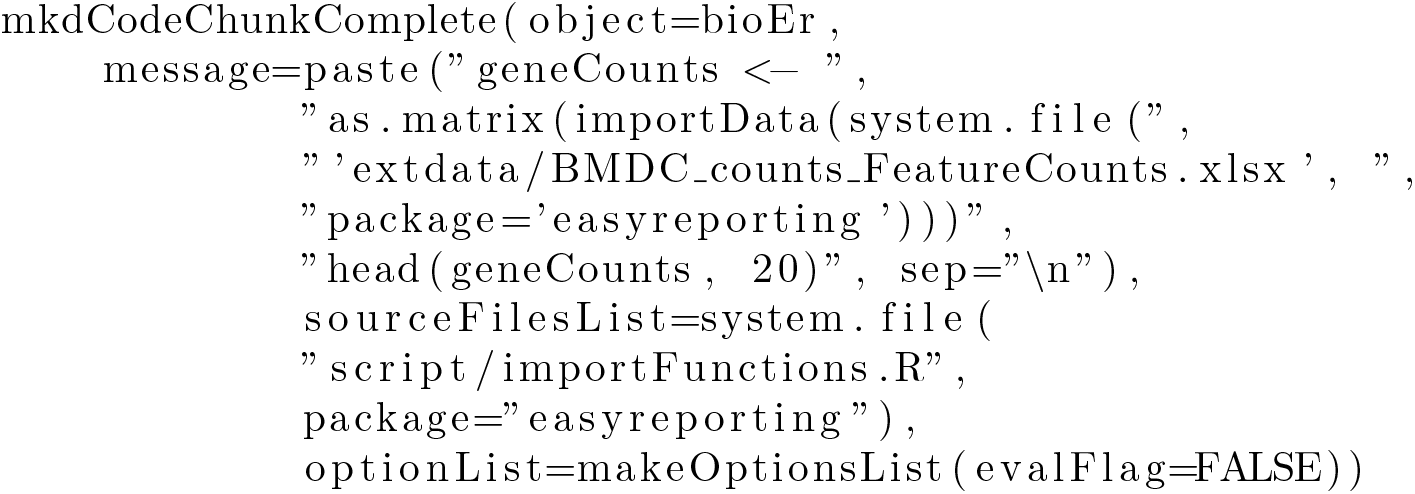
One command chunk construction

Note that the *mkdCodeChunkComplete* allows also to provide specific options for the CC that we are creating. In particular, in this case, if we turn the *evalFlag* to *FALSE*, the code will not be compiled during the final report construction.

It is possible to organize the report in both cases using the *mkdTitle* function. The user has to repeat this operation for each step of the analysis, as shown in the Supplementary File 1. At the end of the process, it is possible to compile the *easyreporting* instance and obtain the analysis report as in Supplementary File 2 in HTML format.

### Implementing automatical tracing functions

This section shows a possible approach on how to encapsulate *easyreporting* methods into third-parties functions to trace the analysis step and execute the code automatically.

First of all, the developer has to write an R function that performs the analysis step of interest (such as the MAedgeRMAPlotEx function for rendering an MA-plot, in our example). Listings 4 shows a simple example of rendering function.

**Listing 4.**
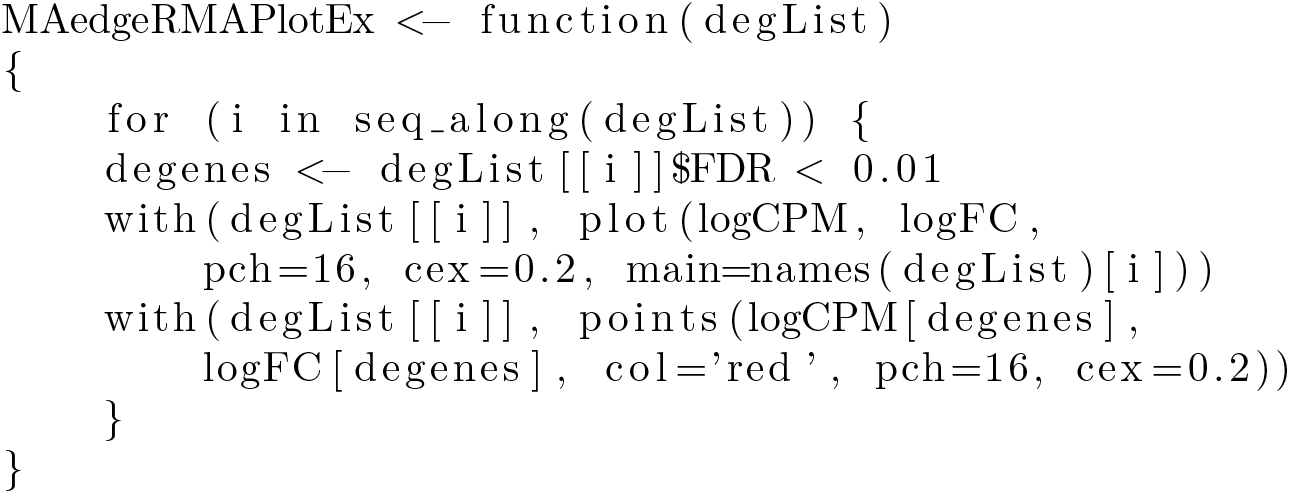
MA-plot rendering function

Note that the developer does not require any extra effort at this stage. Moreover, the rendering function could also be any function available from other packages.

Then, the developer needs also to write a wrapper function (here traceAndPlotMAPlot). The wrapper function should take as input the arguments of the rendering function (here MAedgeRMAPlotEx), and a generic *easyreporting* object (here *er*). Moreover, the wrapper function has to call the *mkdCodeChunkTitledCommented* function of *easyreporting* (where we insert the rendering function call to be traced (MAedgeRMAPlotEx) in the *codeMsg* argument) and the call to rendering function (MAedgeRMAPlotEx). Listings 5 shows the wrapper function of our example.

**Listing 5.**
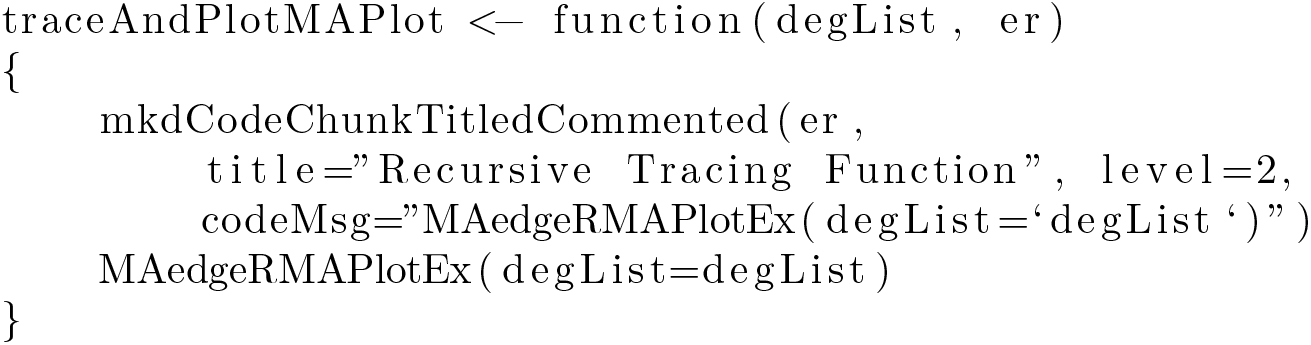
Tracing wrapper function chunk

In this way, the wrapper function allows both to show the result and trace the function in the same step. It is easy to place the *traceAndPlotMAPlot* function-call wherever needed in the main code.

Listing code 5) shows how to use the wrapper function. In particular, we pass as input i) the object required by the rendering function (an edgeR result class in this particular example), ii) an *easyreporting* class instance (here it is the bioEr instantiated in the example).

**Listing 6.**
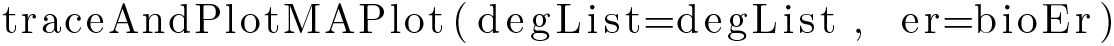
Tracing function call chunk

Note that writing the wrapper function is the only extra effort required to achieve reproducibility.

### easyreporting for GUI implementation

To better illustrate how to incorporate a RR layer into a GUI, *easyreporting* (since version >= 1.3.1 released with Bioconductor 3.13) contains a simple ShinyApp example for plotting a Volcano plot. The command *erGUIVolcano()* allows executing the app and opens the user interface. The user interface allows the user to choose a threshold for the P-value (i.e., the *P-value threshold* for detecting the significant genes in this example) and provide a text area for adding comments. In the interface, there are also two buttons (*Perform Plot* and *Compile Report*) for executing the plot and compiling the report, respectively. The *ui* function provides the code for the user interface. This code does not need to be modified to allow reproducibility. Instead, the back-end of the interface that executes the job has to incorporate a wrapper function (here *traceAndPlotVolcano*), and the call to the wrapper function, respectively. In this example, we also added the report’s initialization. The *server* function provides the code for the back-end interface.

Figure 2 right side shows a schematic representation of the user interface as in the *ui* function. The left side illustrates the back-end as in the *server* function. By using the *Perform Plot* button (red box), the user activates the WF into the server-side, which in turn performs the plot and traces the executed function (the red cascade). In blue is highlighted the argument value and how it is traced through the function cascade.

Additionally, when the user adds its comments (yellow box) to the performed analysis step, the text is passed to the server.

Finally, the user can compile the report using the *Compile Report* button (green box). In this way, the server executes the *compile()* function and produces the HTML report that is automatically showed to the user.

This simple example can be generalized to complex interfaces in order to trace all user interactions.

## Conclusions

*Easyreporting* can be used to support RR in different analysis contexts. However, it is particularly suited for analyzing omics data and developing software/GUIs, as we have shown in this work. Compared to other previously proposed solutions such as [21], that require not negligible commitments by the final user, potentially bringing him/her to renounce to include RR inside the scripts, the implementation of a RR layer with our approach is straightforward. Moreover, it leaves maximum freedom to the developer/analyzer for automatically creating and storing an *rmarkdown* document and providing methods for its compilation and adding comments in natural language.

Although several functionalities are already available, *easyreporting* can still benefit from some extra features such as methods for file editing, graphical representation of the analysis, and data caching. In particular, file editing can be useful for modifying specific CCs, and the graphical representation of the analysis can be useful to provide reports not only readable by third-party users but also graphically visualized as workflows. On the other hand, even though a dedicated data caching infrastructure can offer more manageability and share-ability of the data at the moment, it can be already performed in *easyreporting* by *rmarkdown* CCs option flag.

Finally, thanks to its versatility, *easyreporting* can be ideally included in any well-structured R project and the development of GUIs, helping to fulfill most of the proposed rules in [2]. Moreover, if combined with other tools such as dockers (i.e., *docker4seq* [22]) it helps to create fully reproducible projects. *Easyreporting* naturally complements dockers in terms of reproducibility, allowing both the preserve code lines and user parameters and the computational environments and dependencies.

To conclude, our approach still requires the developer’s effort to implement a RR layer into their software, which makes us imagine possible future works in this area where the code tracing is entirely left to the machine. The Java language provides a well-known example that uses the Aspect-Oriented Programming (AOP) paradigm for the software logging aspects. Unfortunately, this paradigm is still missing in the R language, but possible future approaches in the Reproducible Research area inside R could rely on implementing it, which, combined with *rmarkdown* or similar procedures, can be used to trace ad-hoc tagged functions and to log them into the report file. In such a way, reproducibility could be easier to implement and lesser subject to human errors.

## Supporting information

Supplementary File 1 provides an illustrative example for the creation of an analysis report.

Supplementary File 2 provides the report file obtained using the analysis steps described in the Supplementary File 1.

## Author Contribution

Dario Righelli, Conceptualization, Data curation, Formal analysis, Methodology, Software, Writing – original draft and Claudia Angelini, Conceptualization, Funding acquisition, Methodology, Supervision, Writing – original draft

